# Identification of Key Ischemic Stroke Genes by Computational Systems Biology

**DOI:** 10.1101/2021.10.07.463458

**Authors:** Rongting Yue, Abhishek Dutta

## Abstract

Stroke is one of the leading causes of death in humans. Even if patients survive from stroke, they may suffer sequelae such as disability. Treatment for strokes remains unsatisfactory due to an incomplete understanding of its mechanisms. This study investigates Ischemic Stroke (IS), a primary subtype of stroke, through analyses based on microarray data. Limma (in R)derives differentially expressed genes, and the protein-protein interaction (PPI) network is mapped from the database. Gene co-expression patterns are obtained for clustering gene modules by the Weighted Correlation Network Analysis (WGCNA), and genes with high connectivity in the significantly co-expressed modules are selected as key regulators. Common hubs are identified as Cdkn1a, Nes and Anxa2. Based on our analyses, we hypothesize that these hubs might play a key role in the onset and progression of IS. Result suggests the potential of identifying unexplored key regulators by the systemic method used in this work. Further analyses aim at expanding candidate genes for screening biomarkers for IS, and experimental validation is required on identified potential hubs.

## Introduction

Stroke, as a cerebrovascular disease, takes the third most significant fraction of deaths in the world while also being a leading cause of disability[1]. Moreover, patients who have survived a stroke may still lose independence and require long-term healthcare[2] with great economic burden[3]. As a heritable disease caused by blood blockage in brain artery, IS is a common subtype that makes up 87% of all strokes[4]. However, diagnosis of IS based on clinical brain imaging is not always accessible, and biomarkers based on gene expression profiles are more efficient in identifying strokes[5]. Therefore, the continuous efforts to find out the risk factors of IS are significant, so are the efforts for active prevention, accurate diagnosis and points for therapeutic interventions[6].

Among several stroke models of rodent animals that have been investigated to explore treatments for IS, proximal occlusion of the middle cerebral artery (MCA) via the intraluminal suture technique (so-called filament or suture model) is not only the most frequently used but also the most clinically relevant model that mimics the mechanism of human IS[7], [8]. This model has been introduced as mouse injury model for experimental stroke research[9].

Over the past decade, investigators have utilized gene expression profiles based on data from microarrays to detect key regulators of disease mechanisms, since the profiles have the potential to serve as disease-related biomarkers. Probes in microarrays are capable of measuring thousands of gene expression levels or intensities simultaneously. These high-throughput technologies identify physical and functional relationships between proteins and genes on a large scale. However, mechanisms of IS’s causes and progression relevant to abnormal gene expressions remain largely unclear. Therefore, more comprehensive information about abnormal gene expressions on IS needs to be implemented to intervene stroke process more efficiently. One way to perform gene expression analyses is to find out differentially expressed genes (DEG), which tell abnormal gene up- and down-regulations that are responsible for diseases of interest[10], [11]. The resulting DEGs can be further investigated through enrichment analysis based on different databases. Gene co-expression is another approach that finds out biomarkers for prediagnosis from the view of systems biology, and it has uncovered significantly novel signaling pathways for stroke[12], [13], [14].

In the present study, an analysis based on these two methods was performed to compare gene expressions of mice with Middle Cerebral Artery Occlusion (MCAO)-induced stroke and mice with sham operations. A small group of significantly differentially expressed genes were identified as key regulators. The result has been further filtered by co-expressed gene clusters in the network based on the co-expression information from the correlation between genes and gene expression similarity from network topology level. The core genes in the module were excavated to ascertain the biomarkers highly correlated with the diagnosis of IS. The resulting protein-coding genes Cdkn1a, Nes and Anxa2 were predicted to account for the pathogenesis in stroke process. The next step of this study will focus on expanding hub candidates in geneset and validation through the combination of functional enrichment analysis and other bioinformatics methods to potentially provide new therapeutic targets for the diagnosis and intervention of IS.

## Results

### Differentially Expressed Genes from Microarray

High throughput microarray provides statistics of gene expressions. Gene expression data of GSE35338 and the description file of GPL1261 were downloaded from the NCBI GEO database. After microarray normalization, the differentially expressed genes between the stroke samples and the controls were analyzed. A total of 21746 gene profiles were extracted and converted from GSE35338. Gene expression data was analyzed for three different time points according to description file.

Cut-off criteria of fold-change greater than 1.5 and *P* - values lower than 0.05 was chosen to filter significantly differentially expressed genes in mice with the stroke than the sham surgery. As a result, for up-regulation, 1130 genes, 466 genes and 414 genes were identified for the first day, third day and seventh day, respectively, after treatments. For down-regulated genes, 919 genes, 277 genes and 25 genes were identified, respectively. Venn diagrams indicate that for these three groups after treatment, 139 genes were detected in common for up-regulation, and 3 genes were for down-regulation (Fig.1a and Fig.1b). The common genes were mapped into the STRING database to get PPI network. The interaction confidence of PPI was set to be higher than 0.7. The network was imported to Cytoscape (Fig.2a) and all the disconnected genes were removed for topological analysis. Proteins with degrees more than 10 and stress centrality greater than 1000 were selected as hub proteins (Fig.2b). As a result, 27 proteins in the network were chosen as key regulators for MCAO-induced stroke.

**Fig. 1:**
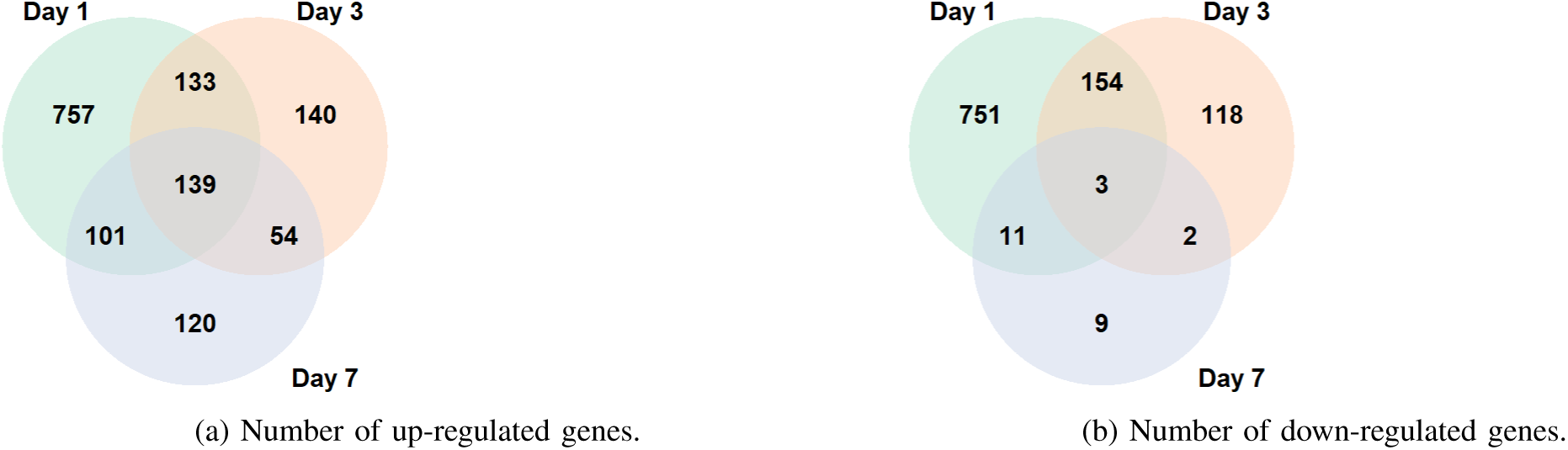
Venn diagrams of up- and down-regulated genes when detecting DEGs with “Limma” package in R. To identify abnormal gene expressions in all three sampling time points, the genes intersected by all time points are chosen.

**Fig. 2:**
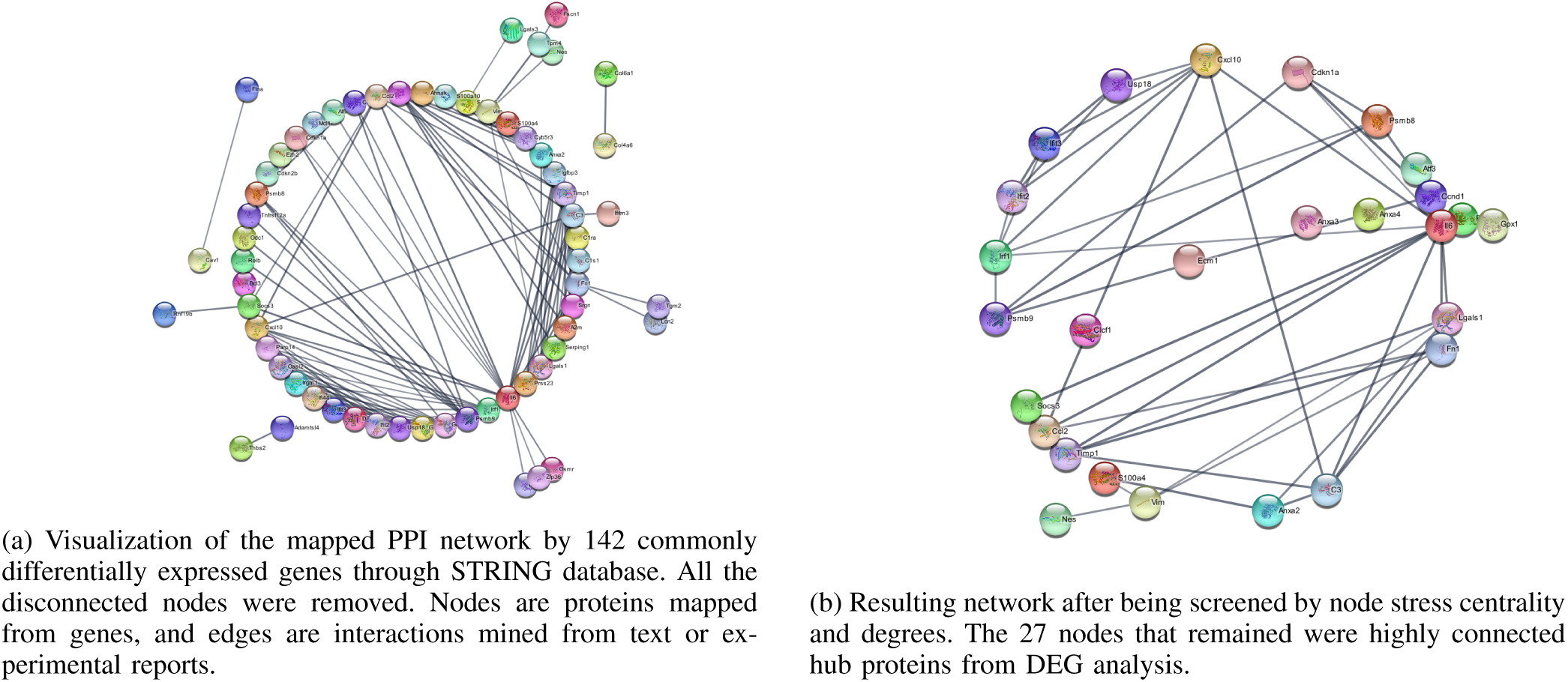
Construction of PPI network by mapping DEGs into STRING database.

### Gene Co-expression

All gene probes (after matching gene expression data with gene symbols from GEO Platform) were re-arranged and filtered according to their mean expression levels. All 21746 genes were ranked in descending order. The first 6,000 genes were used in our work for the subsequent analysis. Since metabolic networks in all organisms have been suggested to be scale-free networks and scale-free network phenomena has been observed in many empirical studies[6], a co-expression network approximated to the scale-free structure was obtained by balancing connectivity and scale-free topology fit index. Gene correlation matrices were calculated and raised to a soft-thresholding power to establish a co-expression network. The scale-free topology fit index fails to go above 0.8 with suitable mean connectivity (Fig.3) for the 21 samples from the dataset. In this case, we go with power 8 that provides mean connectivity of around 165 and median connectivity of around 131, which is reasonable for the trade-off between the scale-free topological index and mean connectivity. This power value reduces the noise when computing correlations of gene expressions.

**Fig. 3:**
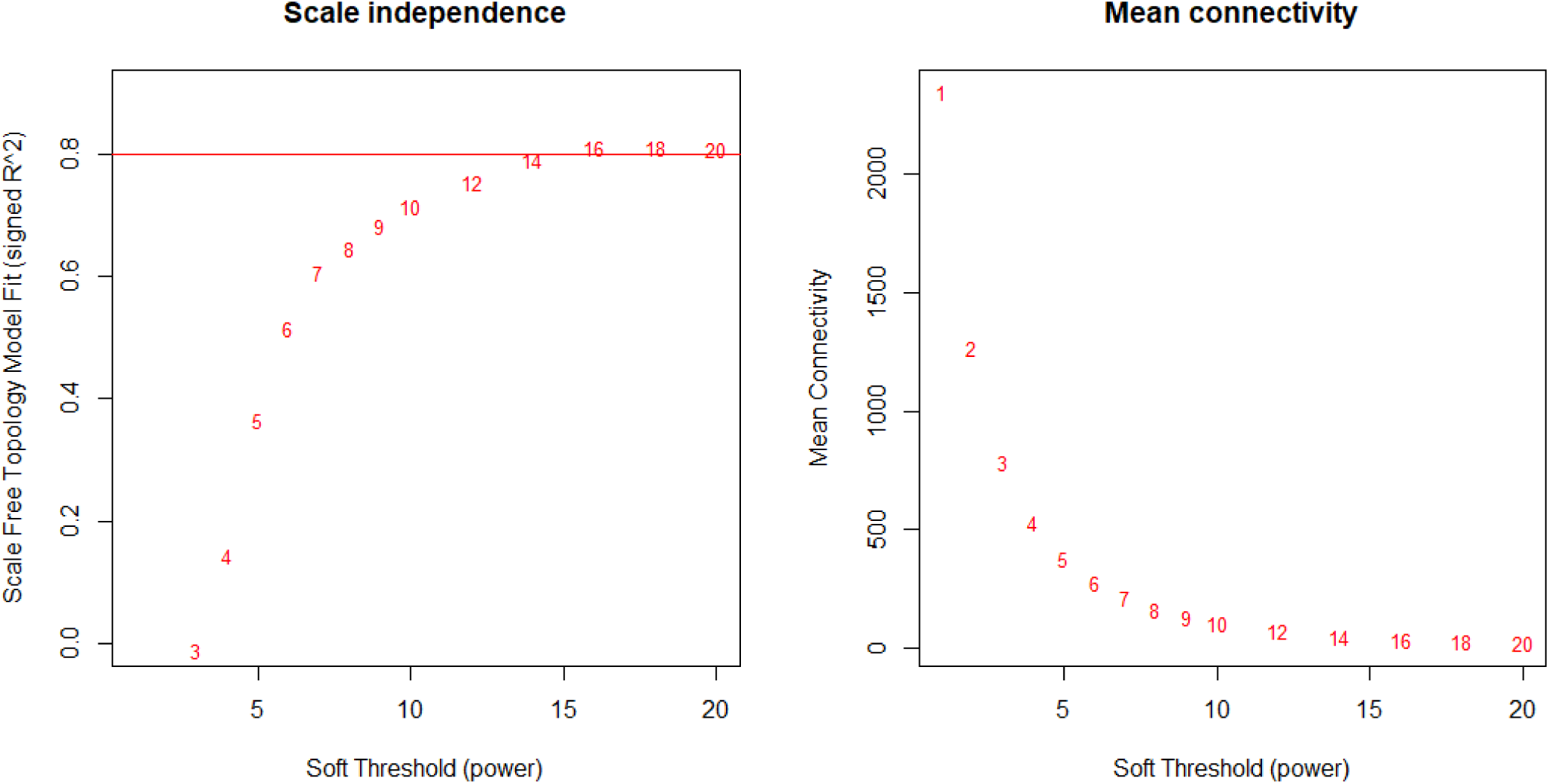
Choice of a soft threshold for gene co-expression clustering based on WGCNA. Scale independence and mean connectivity for different soft-thresholding powers. Higher soft thresholds correspond to lower mean connectivity. The power that makes the scale independence higher than 0.8 can not provide a suitable mean connectivity as shown in the plots.

A hierarchical clustering tree was constructed to identify the gene co-expression network, following the procedures of WGCNA. As shown in Fig.4a, the threshold of cut height (refers to the maximum dissimilarity that qualifies modules for merging) was set to 0.2, such that the modules higher than this height were dissimilar. Genes in the grey module were the ones that have not been assigned to any other modules. A total of 21 modules were identified except for the grey module, and 9 modules were obtained after merging (Fig.4b).

**Fig. 4:**
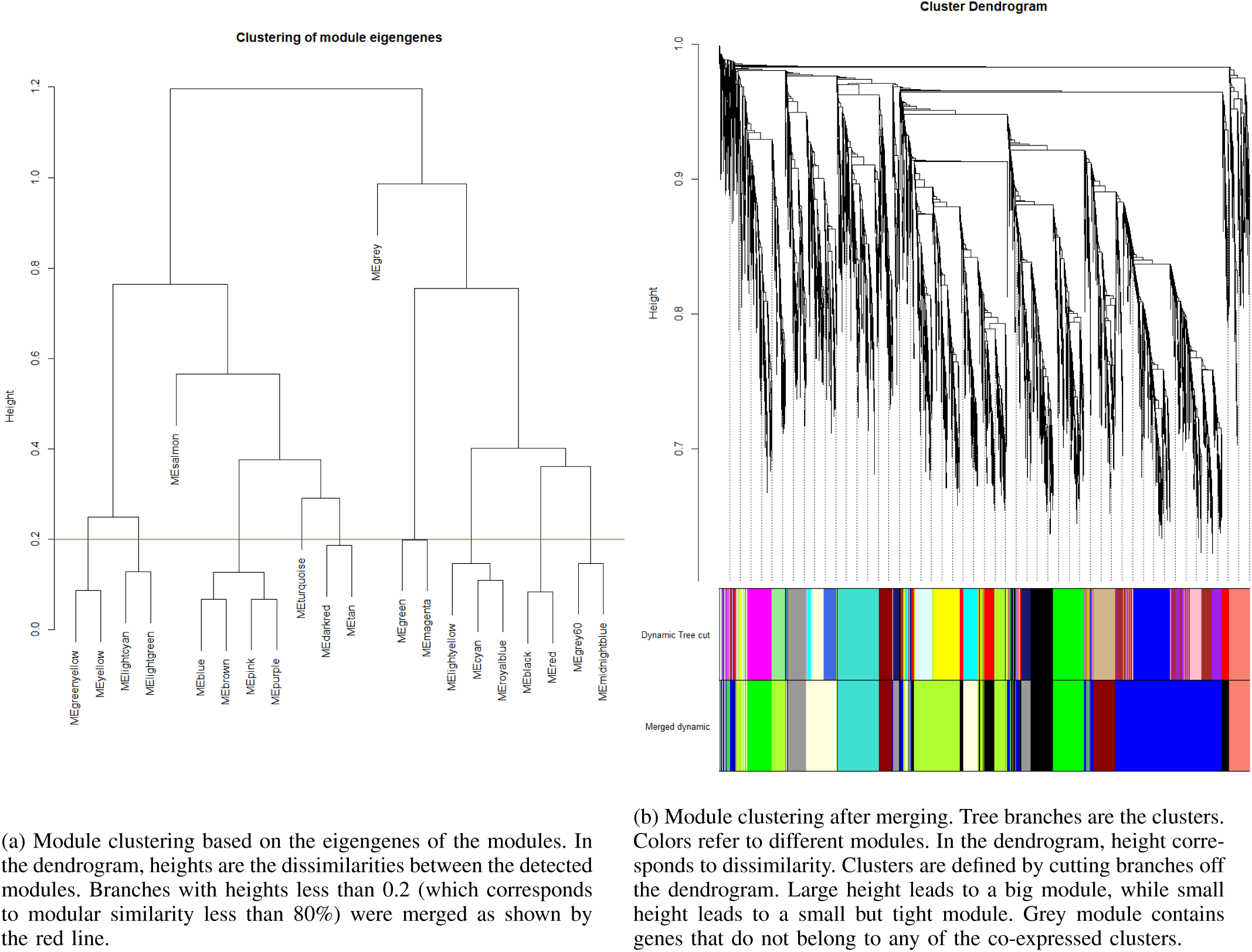
Detection and clustering of the co-expressed modules based on dendrogram.

With mean gene significance across modules (Fig.5a), two modules labeled with “lightyellow” (containing 584 genes) and “greenyellow” (containing 1033 genes) were identified to ascertain modules that were more likely to account for MCAO-induced stroke. Intramodular connectivities were calculated, and the genes with the highest 30% from each module were used (175 genes from the “lightyellow” module and 310 genes from the “greenyellow” module) to select hubs inside these modules. These genes were chosen as the hubs in stroke-related gene modules. As a result, Venn diagram in Fig.5b shows 3 genes Cdkn1a, Nes and Anxa2 in common for gene sets from DEG and co-expression. The resulting genes are biologically and statistically meaningful, since their expressions are varied significantly, according to the microarray data, and they are drawn from the hub candidates in two significant gene co-expression modules in organisms with stroke. These 3 common hub genes are hypothesized to play an important role in the onset and progression of IS.

**Fig. 5:**
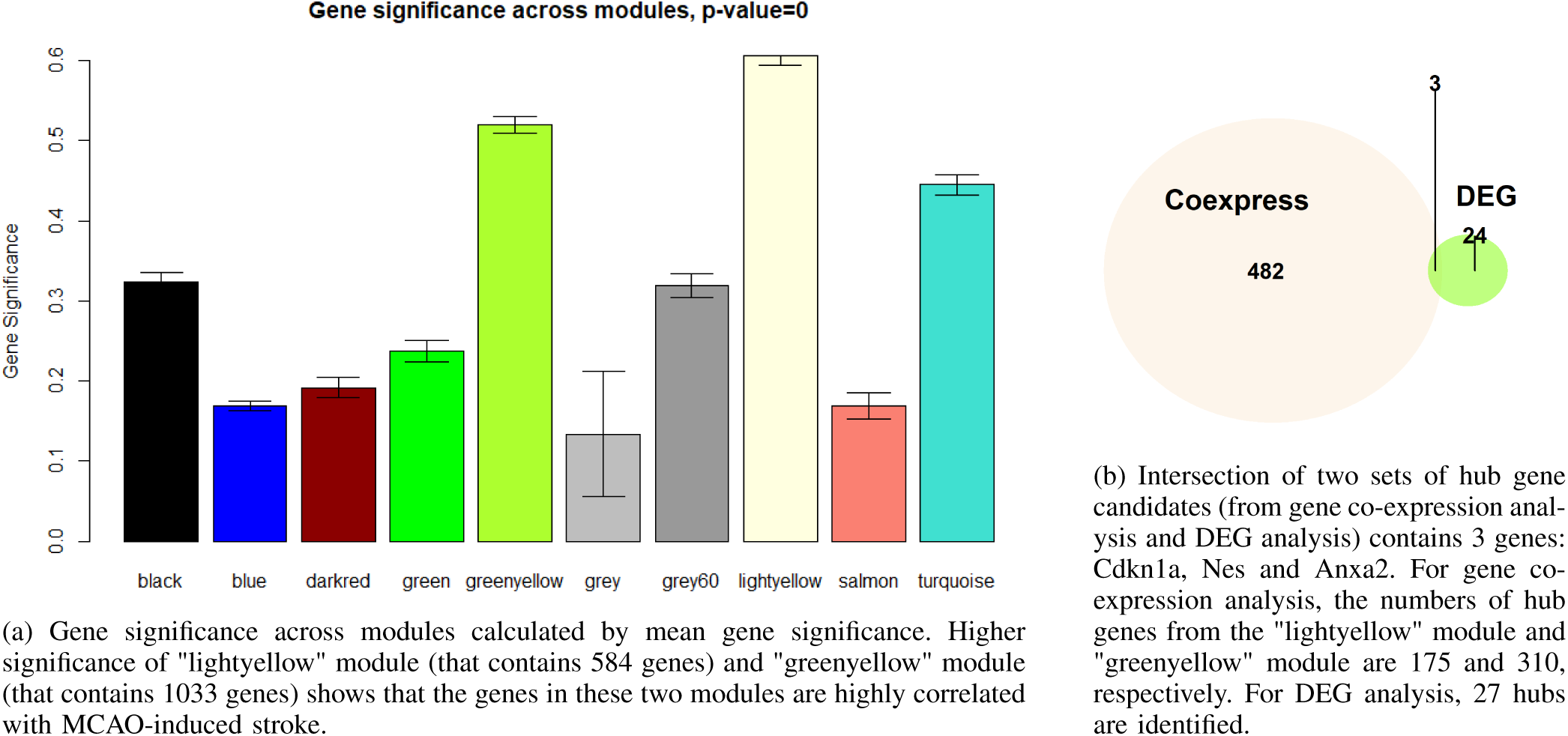
Analyses of the selected modules based on gene co-expression.

## Discussion

IS is a major subtype of stroke with an unclear predominant etiology. The differentially expressed gene products may be the biomarker in the processes of brain damage after stroke, including axon guidance and apoptotic response to cerebral ischemia[15], [10]. More risk factors of IS remain to be fully elaborated for therapeutic interventions[6].

Analyses based on microarray data have gained increasing attention because they acquire high throughput gene expression data simultaneously and provide us comprehensive knowledge of variation in gene expression to increase our understanding of normal and diseased states[16]. In this study, we use microarray data of mice model with MCAO-induced stroke since it is one of the most frequently used ones for the capability to mimic the mechanism of human IS[7], [8]. We analyzed the genetic profiles of reactive and quiescent astrocytes in young adult mouse brains with MCAO-induced stroke from GEO database. Since the methodologies of identifying genetic differential expressions and co-expression are highly efficient for targeting disease-related genes and proteins[17], they provide potential clinical diagnoses, prognostic biomarkers and therapeutic targets for acute ischemic stroke via the identification of DEGs and co-expressed genes.

We first detected altered gene expressions from microarray. DEG analysis showed there were 139 genes and 3 genes in common for up- and down-regulation, respectively. These genes were detected significantly differentially expressed in mice with MCAO-induced stroke. PPI network was constructed by mapping DEGs into STRING database, and a total of 27 hub proteins were identified through topological parameters. The hubs were key regulators that might play an important role in the process of IS. The next phase of this study involved detecting gene co-expression patterns by WGCNA based on an unsupervised manner. Under the assumption that all genes are connected, WGCNA quantifies gene connection strength with the expression correlations. 9 modules were detected (besides the grey module), and 2 of them had higher mean gene significance, which indicated that genes within them were more likely to account for IS. Further in this study, the resulting protein-coding genes Cdkn1a (Cyclin Dependent Kinase Inhibitor 1A), Nes (Nestin) and Anxa2 (Annexin A2) intersected by two methods above were identified as the common hubs, which were predicted to be biologically and statistically meaningful. We hypothesized that the common hubs are most likely to play an important role in the onset and progression of IS based on our analyses.

Cdkn1a encodes cyclin-dependent kinase inhibitor that regulates the cell cycle progression at G1. This gene has been widely investigated as a tumor suppressor for tumor progression and prevention. Though there are not many studies that reveal the direct correlation between Cdkn1a and IS or brain injury, indirect evidences can be found that Cdkn1a is the target gene of p53[18], and inhibition of p53 pathway intervenes the process of neurogenesis after IS[19]. Single-nucleotide polymorphisms (SNPs) in the human Anxa2 gene have been associated with the increased risk of stroke[20]. Also, Anxa2 has been studied to connect stroke with several SNPs in the Endothelin 1 gene (on chromosome 6)[21]. In the rat thrombosis models, the potential clinical utility of recombinant Anxa2 protein in IS has caught the attention of researchers [22]. Nes is a protein-coding gene in Homo sapiens and it encodes Nestin protein. Nestin is originally introduced in neuroepithelial stem cells and defines intermediate filament (IF) protein in brains[23]. Recent evidence suggests that inflammation is involved in the pathogenic progression of ischemic stroke[24], [3]. Nestin is predicted to serve as a context-dependent marker of activated microglia/macrophages, and microglial expression of Nestin increases in response to neuroinflammation[25]. In atherosclerosis-related research, Nestin cells have been proven to direct inflammatory cell migration during chronic inflammation[26]. Based on experiments, recent studies have validated that Nes expression is closely relevant to ischemic brain[27], [28], confirming our results obtained by more efficient, comprehensive and generalizable computational systems biology methods grounded in rigorous statistical analyses of microarray differential gene expression data. The key to developing effective and clinically applicable treatment methodologies is a better understanding of the pathophysiology of the disease. Structural hubs in molecular interaction networks potentially provide the therapeutic targets that intervene in the causes or progressions of diseases. With the interaction of gene expression products on the molecular level, more potential biomarkers and therapeutic targets can be identified.

Further studies include the expansion of geneset that account for IS by the approaches used in this work. Missing data occurs in microarray during experiments, and it results in an adverse effect on downstream analysis, which requires imputation algorithms to make use of available information to impute missing values[29]. Our study used the intersection of DEGs from all time points, and this might be inaccurate when the measurement of gene expression data is missing, which leads to fewer gene candidates for the screening process. There exists a potential to expand gene candidates and find drug modules that combine target gene expression products to intervene the process of IS. In a co-expression network, important modules for the disease are selected through module significance. The unselected modules may also contain genes that contribute to IS, but they are ignored. The genes with the highest connectivity in those less significant modules may be drawn to expand the co-expressed gene candidates. Also, the co-expression network does not offer causality of gene regulation since the undirected edges lose some regulatory information (e.g., the direction of suppression/activation). Besides, experimental validation of functions of unexplored genes is required to test if they contribute to IS mechanism. Also, drug molecules that bind with these disease-related omics molecules should be detected for therapeutic purposes, since specific antiinflammatory interventions are beneficial either in prevention or acute treatment. Through drug-drug interactions, drugs have the potential to be re-purposed or combined to improve current therapeutic performance for IS.

## Materials and Methods

### Data and Pre-processing

Microarray collects the expression levels of different genes in the form of fluorescent intensity. Gene expression profiles of young adult male mice in GSE35338 (Seris matrix of Mus musculus data) come from platform GPL1261 in Gene Expression Omnibus (GEO) database. Samples with treatment labels “sham surgery” (controls) and “MCAO-induced stroke” (cases) are selected for analyses. Data is sampled from the purified reactive astrocytes after these treatments.

The probe-level data in Affymetrix probe result file were converted into expression measures and normalized to account for technical variation by Robust Multichip Average algorithm[30], since this technique does not utilize mismatched spots in the microarray. This algorithm is made up of three steps: background correction, normalization and summarization[30]. More specifically, background correction is performed based on the probe signal *S* and background noise *BG*, and the model observed perfect match *PM* of RNA probes

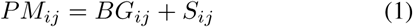

where *BG* ∼ *N*(*µ, σ*), *S* ∼ *Exp*(*r*), (*BG, S >* 0, *BG* and *S* are independent), *PM*_*ij*_ refers to the observed expression profiles from the *j*th probe on array *i*. The purpose of this step is to derive a transformation function *B*, such that

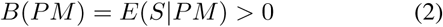

When substituting *BG* = *PM* − *S*, the joint probability density function of *S* and *BG* is

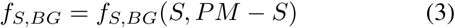

With total probability theorem, the conditional probability density function of signal intensity *S* given perfect match *PM* is[31]

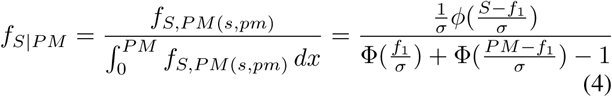

where *f*_1_ = *pm* − *µ* − *rσ*^2^, *ϕ* is the probability density function (PDF) and Φ is the cumulative distribution function of standard normal distribution. Then the background adjusted PM given measurement is[31]

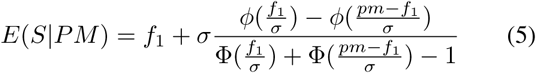

The unknown parameters *µ, σ* and *C* for cumulative density function of *BG* and *S* are estimated through non-parametric kernel estimation[30].

Assume *n* probe values for gene *x* are *x*_1_, *x*_2_, …, *x*_*n*_ that are *i*.*i*.*d*, note their PDF as *f*, then kernel density estimation is

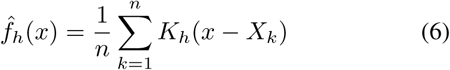

where 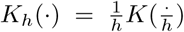 is a scaled kernel, positive *h* is bandwidth used as a smoothing factor, and *K*(·) is the kernel function that is non-negative and *E*[*K*(·)] = 0 and ∫ *K*(·) *dx* = 1 (e.g., Gaussian kernel for microarray data[32] is 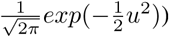

Then the quantile normalization is performed to make each gene expression distribution identical in statistical properties, such that expression profiles in each sample have identical quartiles and the order of gene expressions within each sample is reserved. The last step, summarization, uses Tukey Median Polish method to reduce noise in an array. The smoothing operations are needed to eliminate the column and row effect in the gene expression matrix. Assume the rows and columns in expression matrix are *n* samples and *m* genes, respectively, note the expression profile of sample *i* and gene*j* as *X*_*ij*_, then procedures of Tukey Median Polish method in each iteration are listed in Algorithm 1. Data after normalization are almost within the same range. The mean expression levels are used for the repeated probes. (When performing dendrogram clustering based on Euclidean distance and Complete-linkage, only a few outliers are present.)

#### Algorithm 1: Tukey Median Polish

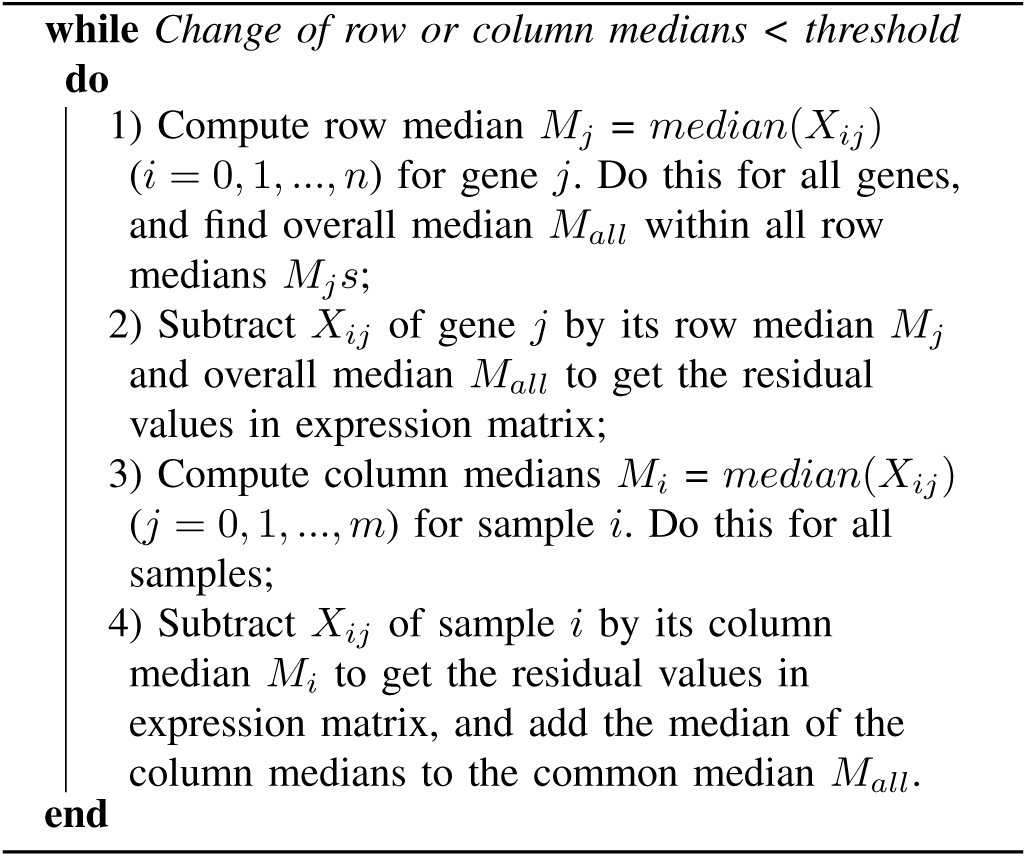

### Identify the Differentially Expressed Genes

The statistics of gene expression profiles are analyzed by “Limma” package[11] (in R) to find out the Differentially Expressed Genes (DEG). Data is grouped according to the first day, third day and seventh day, respectively, after treatment. The empirical Bayes moderation is utilized in the algorithm, since an inverse Chi-square prior is estimated from data[11]. Then data is fitted into multiple linear models, and the moderated t-statistics are computed[11]. The following steps are adapted from the DEG analysis based on “Limma”.

A linear model from several microarray for particular gene *k* is assumed to have mean *E*[*y*_*k*_] = *XC*_*k*_ and variance 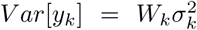, where *y*_*k*_ is the obtained expression profile of gene *k* (i.e., *y*_*k*_ is the *k*th row of expression matrix), *X* is the design matrix that explores differences between target samples (2 columns are sample traits with MCAO-induced stroke and sham surgery, and rows are different samples), *C*_*k*_ is coefficient vector, *W*_*k*_ is weight matrix, and 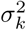 is the variance of expression profiles of a specific *k*[11]. Nonzero values on the diagonal refer that the measurement data is present. The contrast matrix *F* makes the pairwise comparisons and specifies the hypothesis needed to be tested, and it measures the change between cases and controls. Contrasts of coefficients is defined as *ϵ*_*k*_ = *F*^*T*^ *C*_*k*_.

Linear models are fitted to gene expression data to obtain coefficient estimator *Ĉk*, sample variance estimator 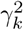 and estimated covariance

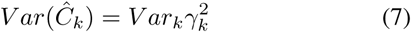

where *V ar*_*k*_ is a positive definite matrix independent of 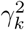. The contrast estimator 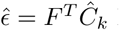 has estimated covariance

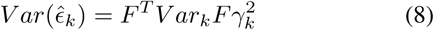

The contrast estimator is nearly normal with mean *ϵ*_*k*_ and covariance matrix 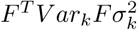. Note *d*_*kj*_ as the *j*th diagonal element of *F*^*T*^ *V ar*_*k*_*F*, then given *ϵ*_*kj*_ and 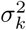, the individual contrast estimator 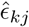 of gene *g* follows normal distribution:

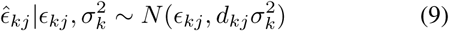

Given 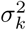, sample variance estimator 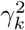 follows Chi-square distribution with a scalar 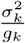, where *g*_*k*_ is residual degrees of freedom for a linear model of the specific gene *k*.

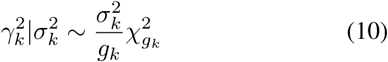

The inverse Chi-square prior that derives variances 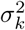 among genes is estimated with prior estimator 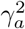 and *g*_*a*_ degrees of freedom. (*g*_*a*_ and 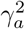 and are estimated from observed/residual sample variances 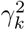 that follows nearly Chi-square distribution. These two parameters are obtained through setting the empirical to the expected values for mean and variance of the finite and nearly normal log 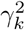)

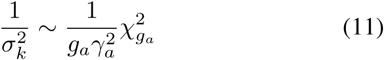

Given prior observation with variance *d*_0*j*_ at *j*th time point, log-fold-change of DEGs follows distribution 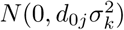, where *d*_0*j*_ is estimated from moderated t-statistics *t*_*kj*_ with knowledge of *g*_*a*_ and *p*_*j*_ (i.e., probability of non-zero *ϵ*_*kj*_ or valid portion of DEGs).

The ordinary *t*-statistic is

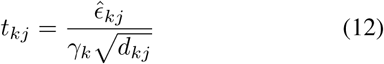

where *d*_*kj*_ is the *j*th diagonal element of *F*^*T*^ *V ar*_*k*_*F* and it is the unscaled standard deviation, and the moderated t-statistics *t*_*kj*_ with added degrees of freedom *g*_*k*_ + *g*_*a*_ has the form

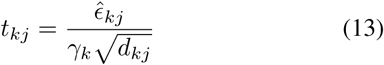

that sample variance 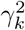 is replaced by posterior variance 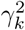, which is

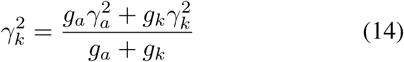

From the statistical result, 1.5 fold-change of genes expression (corresponds to 0.58 in logarithmic scale) in a Student’s t-test across replicas helps select the significantly variant genes[33]. Note that positive fold-change refers to up-regulation of genes and negative value refers to down-regulation. *P* -value that describes the statistical certainty of genome-wide significance often uses 0.05 as the threshold such that the test is believed to be not by chance with 95% certainty[34]. Gene expression profiles on microarray are sampled at first day, third day and seventh day, respectively, after treatment, according to the description file. Therefore, the up-regulated and down-regulated genes should be analyzed separately. Enrichment analyses can further map these DEGs into KEGG database to detect abnormal gene expression according to signaling pathways.

The resulting DEGs are mapped into the STRING database. After setting the confidence level of the interactions, a set of Protein to Protein Interactions (PPIs) are downloaded from the database directly. Only a few genes are responsible for regulation. Since gene regulations are many-to-many, network structure offers an excellent visualization to describe their relations. PPI network is visualized in “Cytoscape” and analyzed with toolbox “NetworkAnalyzer”. Hub proteins that act as key regulators can account for most of the disease mechanisms and can be identified with topological parameters[35]. Stress centrality can be a criterion to select the “hub”, since it describes the capability of a protein to connect nodes as a communication center in a protein network[36], which is computed as follows:

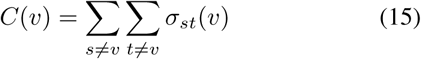

where *σ* denotes the number of shortest paths containing edge *v, s* and *t* are nodes that are different from the given node *v*. Another criterion used to get hub proteins is the degree. Degree records the number of nodes connected directly to a given node. High degree value indicates that the nodes are likely to be hubs in a certain network[36]. The resulting DEGs are too many, although the burden can be partially eased by adjusting thresholds, e.g., larger fold-change. Also, in DEG analysis, comparisons between samples are performed for all time points, which means the complexity and the computational load increases when data is sampled from several time points. And no information is available on edges in the PPI network.

### Clustering Based on Gene Co-expression

To not analyze gene expressions according to timestamps, WGCNA[37] (in R) can be incorporated to screen real hubs[38]. It provides an unsupervised analysis method that clusters genes based on their expression profiles. A co-expression network is established that approximates the scale-free structure by properly raising gene expression correlations to a soft-thresholding power, enlarging the differences between correlations. It identifies the highly co-expressed gene sets in which those genes have identical expression patterns. For example, if the samples are all control (or case), all the co-expressed genes should be up-regulated or down-regulated within those samples. Weights in the network annotate correlations. A biological scale-free network has few hub nodes with plenty number of connections, and the remaining nodes have fewer connections[39]. Two advantages of the scale-free network are that a new node will be more likely to be connected to the hubs, and the network is robust to accidental failures. Besides, the clustering approach based on gene co-expression outperforms principal component analysis when dealing with high-dimensional gene expression data, since it does not focus on the dimensions of data with the highest variation[6].

With gene expression profiles, Pearson’s correlation between a pair of gene expression profiles *x*_*i*_, *x*_*j*_ can be calculated:

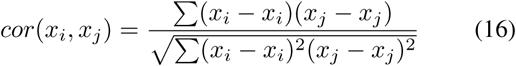

According to WGCNA, the co-expression similarity is derived as the absolute value of this correlation:

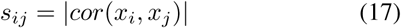

Adjacency matrix describes connection strength between gene expression profiles in the proposed weighted network and it is calculated by raising the co-expression similarity to a power:

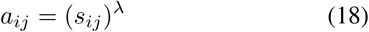

where soft-thresholding power *λ* is used to reduce the noise of the correlations in the adjacency matrix and used in co-expression modules. Key concept in WGCNA is connectivity that describes the relative significance of a gene in network[6]. Connectivity measurement for each gene is the sum of the connection strengths between that gene and all the other genes in the network, and the mathematical representation is 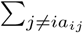 for the *i*−th gene, *a*_*ij*_ is the adjacency of gene *i* and *j*. Mean connectivity *C*_*m*_, also refers to network density[40], is then calculated as follows:

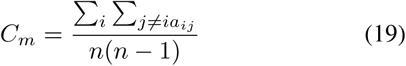

The higher power makes the network closer to a scale-free structure, but it results in lower mean connectivity. Generally, a kink appears in the scale independence as soft threshold changes to reserve mean connectivity, since the scale-free topology fit will not improve much after the kink[41]. A larger scale-free topology fit related to connectivity refers to a high signed *R*^2^, indicates that the model explains most of the variability of the response data around its mean. A proper *λ* can then be chosen to balance the scale-free topological index and mean connectivity. Since Low gene expressions are usually regarded as noise, the normalized gene expression profiles are filtered by mean expression (or variance) for robustness. A topological overlap matrix (TOM) is then calculated to measure the topological overlap of two genes *i* and *j* as follows:

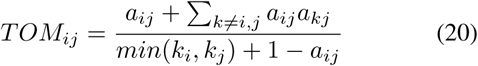

where *k*_*i*_ = Σ*j a*_*ij*_. The similarity measures in TOM take the range [0,1] and the dissimilarity is then derived by subtracting similarity from 1. Then, an agglomerative hierarchical clustering strategy is applied using the TOM-based dissimilarities as the merging distances of the branches, and these branches are merged with a bottom-up strategy through dynamic hybrid cut algorithm[42]. Member quantity *M* within a given cluster core is as follows:

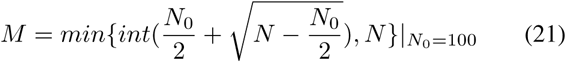

where *N*_0_ is the minimum cluster size (chosen to be *N*_0_ = 100) and *N* is the total number of objects in the cluster. Small clusters are merged into the nearest larger clusters within the same branches in dendrogram based on average dissimilarities, and tiny clusters that fail to meet the size requirement can be assigned to the closest basic cluster even if they are not on the same branch. The other unlabelled objects remain to be individuals.

A cut height (maximum joining height) of 0.2 is set to fuse modules with 80% similarity and reverse number of clusters as much as possible. Scatters of cluster cores describe the average of all pairwise dissimilarities of the elements between each two of the clusters. A maximum core scatter, based on the 5^*th*^ percentile of joining heights, is used to exclude objects which are far from a cluster, even if they belong to the same branch of the dendrogram. A minimum cluster gap, based on cut height and the 5^*th*^ percentile of joining heights, is used to separate clusters from their surroundings.

Eigen-genes *E*, which are the unique orthonormal super-positions of the genes[43], are used as the representatives to tell differences between one gene module and another by taking the first principal component as a summary of that module, after performing singular value decomposition on the expression matrix. The computation is as follows:

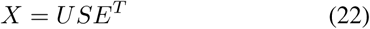

where *X* comprises normalized gene expression profiles, *U* is the matrix with columns as eigenvectors of *XX*^*T*^, and *S* is a matrix with singular values. Modular similarity is calculated by the correlation between the module eigengenes. Modules whose eigengenes are highly correlated are merged. Gene significance is measured by the absolute correlation between the trait (stroke) and the expression profile measures. The modular significance is measured by mean gene significance across modules. A set of significant genes, with the highest intramodular connectivities in the module, are selected as hubs in that module.

Note hub DEGs as set *A*, and the co-expressed genes identified above as *B*, then the intersection geneset *C* can be derived

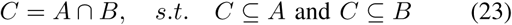

where the intersection of this co-expressed gene set and DEGs are the target genes that are not only biologically meaningful[42], but also statistically meaningful to stroke. The modules that are close to stroke are identified till this step. However, a single module contains hundreds of genes, and the most important genes have to be detected. The correlations between genes and module and between phenotype (stroke) and genes are analyzed jointly to solve the aforementioned issue. These genes are not only highly related to their clustering modules but also highly related to IS.

